# Micro- and macroevolutionary trade-offs in plant-feeding insects

**DOI:** 10.1101/040311

**Authors:** Daniel A. Peterson, Nate B. Hardy, Benjamin B. Normark

## Abstract

A long-standing hypothesis asserts that plant-feeding insects specialize on particular host plants because of negative interactions (trade-offs) between adaptations to alternative hosts, yet empirical evidence for such trade-offs is scarce. Most studies have looked for microevolutionary performance trade-offs within insect species, but host-use could also be constrained by macroevolutionary trade-offs caused by epistasis and historical contingency. On the other hand, evolutionary interactions between adaptations to diverse hosts could be neutral or positive rather than negative. Here we used a phylogenetic approach to estimate the micro-and macroevolutionary correlations between use of alternative host plants within two major orders of plant-feeding insects: Lepidoptera (caterpillars) and Hemiptera (true bugs). Across 1604 caterpillar species, we found both positive and negative pairwise correlations between use of diverse host taxa, with overall network patterns suggesting that different host-use constraints act over micro-and macroevolutionary timescales. In contrast, host-use patterns of 955 true bug species revealed uniformly positive correlations between presence on the same host taxa over both timescales. The lack of consistent patterns across timescales and insect orders indicates that host-use trade-offs are historically contingent rather than universal constraints. Moreover, we observed few negative correlations overall despite the wide taxonomic and ecological diversity of the focal host taxa, suggesting that positive interactions between host-use adaptations, not trade-offs, dominate the long-term evolution of host use in plant-feeding insects.

## Introduction

Most plant-feeding insects are ecological specialists restricted to a small number of host-plant species (Forister et al. 2015). The prevalence of specialization is surprising given the advantages of being a generalist (including greater resource and refuge availability), and many researchers have therefore suggested that the evolution of generalism must be constrained (Futuyma and Moreno 1988; Futuyma et al. 1995; Scriber 2010). This constraint is usually imagined as a trade-off between adaptations to alternative hosts, whereby an increase in performance on one host comes at the cost of decreased performance on another host (Agrawal et al. 2010; Forister et al. 2012). Such trade-offs are crucial elements of most theoretical models of the evolution of specialization (Ravigné et al. 2009; Nurmi and Parvinen 2011; Remold 2012), and are often assumed to arise as consequences of the genetic architecture of host-use. One frequently invoked genetic model involves antagonistic pleiotropy, in which distinct alleles at a single locus have opposite fitness effects on alternative hosts (Futuyma and Moreno 1988; Scheirs et al. 2005; Scriber 2010; Gompert et al. 2015). For example, slightly different enzymes might be most efficient at detoxifying each plant species' unique secondary compounds (e.g. Li et al. 2003). Despite the intuitive appeal of antagonistic pleiotropy, however, empirical studies have generally failed to find evidence for negative genetic correlations between performance on alternative hosts within insect species (Futuyma 2008; Forister et al. 2012; Gompert et al. 2015). Nevertheless, antagonistic pleiotropy may be difficult to detect within species because its effects can be obscured by segregating fitness variation at non-host-specific loci (Joshi and Thompson 1995). Moreover, genetic variation for use of novel hosts is often absent within a single population (Futuyma et al. 1995) and host-use is phylogenetically conserved in many insect groups (Futuyma and Agrawal 2009). We therefore cannot rule out the possibility that historical antagonistic pleiotropy drove the evolution of specialization in ancestral lineages of plant-feeding insects.

Although the prevalence of host-use specialization is often attributed to adaptive trade-offs, some theoretical models suggest that specialization can evolve even when adaptations to one host do not decrease performance on other hosts. Most insect species can choose which host plant they will feed on, so evolutionary feedback between the evolution of host choice and host performance could drive behavioral specialization (Ravigné et al. 2009; Nurmi and Parvinen 2011). For example, if a particular adaptation increases fitness on one host more than on another, individuals may evolve to feed preferentially on the host that gives them higher fitness (Fry 1996). If a non-preferred host is rarely used, selection for performance on that host will be weak, and mutation and genetic drift may eliminate the genetic tools required to use it (Whitlock 1996). In general, over long timescales, the selective environment will shape a lineage's genome, and epistatic interactions between new mutations and their genetic background will determine whether adaptations to novel hosts are possible (Weinreich et al. 2005; Remold 2012). We therefore expect that the evolution of host use is constrained by historical contingency and the complexity of genetic interactions. In fact, the importance of historical contingency and epistasis for the evolution of specialization has been demonstrated empirically by experimental evolution in microbial systems: trade-offs between environments can appear after significant periods of cost-free adaptation (Satterwhite and Cooper 2015) and realized trade-offs can differ between replicate lineages (Rodriguez-Verdugo et al. 2014). On a rugged adaptive landscape, evolutionary trajectories to alternative resource-use strategies may be mutually exclusive, and the direction taken by each lineage can depend on stochastic factors like mutation order (Elena and Lenski 2003).

The adaptive landscape is likely rugged for plant-feeding insects because their interactions with hosts are genetically complex (Ali and Agrawal 2012; Barrett and Heil 2012; Remold 2012). If historical contingency and epistasis constrain the evolution of host-use, adaptations to one set of hosts may reduce the probability of adapting to another set of hosts, driving specialization over long evolutionary timescales. Analagous macroevolutionary trade-offs have been described in plants; alternative defensive strategies tend to be negatively correlated over plant evolutionary history (Campbell and Kessler 2013; Johnson et al. 2014). It remains unknown, however, whether the diversification of host plant defenses has created trade-offs for plant-feeding insects.

Although trade-offs could arise from either genetic architecture or historical contingency, each of these mechanisms could instead produce positive interactions between use of distinct hosts. A single mutation might improve performance on multiple hosts, for instance by improving an effector protein that inhibits a defensive pathway conserved across multiple plant species (Barrett and Heil 2012). Similarly, the appearance of a new enzyme class could create short-term trade-offs as the enzyme is calibrated to different hosts, but long-term performance benefits across multiple hosts after gene duplication (e.g. cytochrome P450 monooxygenases; Li et al. 2003). It is also possible that the genetic factors affecting performance on alternative hosts are independent, experiencing purely neutral interactions on both micro-and macroevolutionary timescales.

One way to investigate the importance of evolutionary interactions between traits is to map the traits onto empirical phylogenies of extant species and ask whether the traits are correlated over the evolutionary history of the focal group (Maddison and FitzJohn 2015). Negative correlations across species suggest trade-offs (Shoval et al. 2012), although correlations alone cannot distinguish between mechanistic constraints and associations shaped by selection pressure (Agrawal et al. 2010). However, recently developed statistical methods allow the partitioning of correlations between species traits into phylogenetic and residual components (Hadfield and Nakagawa 2010). Macroevolutionary interactions driven by historical contingency in ancestral lineages should be apparent in correlations between traits over phylogenetic timescales, while microevolutionary interactions should be captured by residual variation - the evolution that has happened independent of the species' shared ancestry. Phylogenetic analyses therefore allow characterization of positive, negative, and neutral interactions between traits over both short and long evolutionary timescales (Figure 1).

**Fig. 1.**
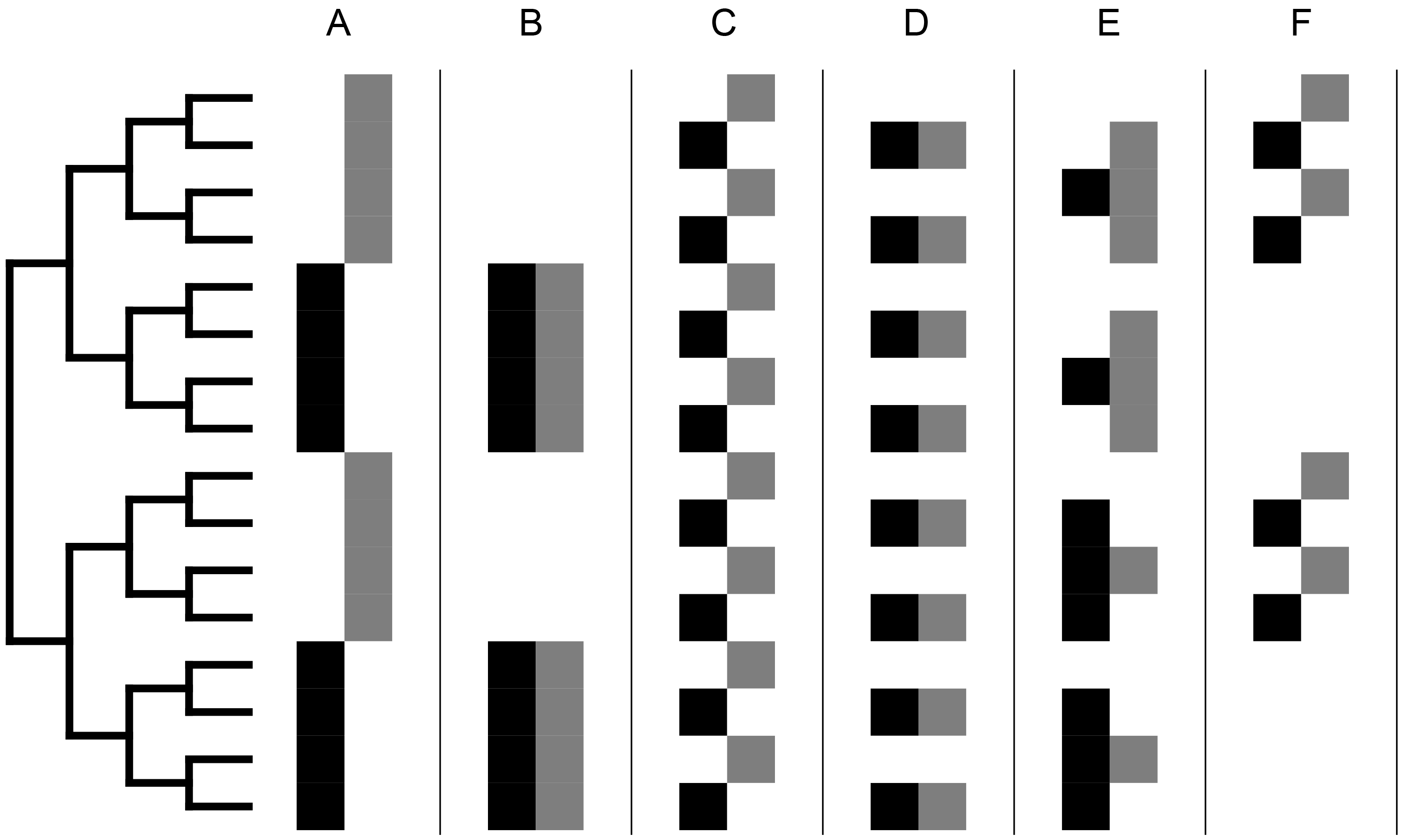
Phylogenetic and residual evolutionary correlations between traits. Hypothetical scenarios of evolutionary correlation between herbivore presence on two hosts: (A) negative phylogenetic correlation, (B) positive phylogenetic correlation, (C) negative residual correlation, (D) positive residual correlation, (E) negative phylogenetic and positive residual correlations, (F) positive phylogenetic and negative residual correlations. In each example, black squares on the left indicate which species in the herbivore phylogeny are present on host 1, and gray squares on the right indicate which species are present on host 2.

Here we used phylogenetic methods to investigate interactions between adaptations to diverse host taxa over micro-and macroevolutionary timescales in two orders of plant-feeding insects: Lepidoptera (caterpillars) and Hemiptera (true bugs). Using digitized insect collection records from North America, we estimated pairwise evolutionary correlations between presence on common host-plant orders across hundreds of species in each insect order. We then combined the pairwise correlations into network graphs, revealing overall patterns of host-use evolution in each insect order. We expected that use of the focal hosts would be mostly negatively correlated or clustered into discrete functional groups if specialization in plant-feeding insects is driven by widespread trade-offs between adaptations to different hosts. A distinction between micro-and macroevolutionary trade-offs could be made by asking whether the negative correlations appeared in the insects' residual or phylogenetic host-use variation. On the other hand, if specialization is not caused by trade-offs between adaptations to alternative hosts, we expected that correlations between host-use traits would be neutral or positive, with little overall network structure.

## Materials and Methods

**Data Collection.** Lepidopteran host-use data were downloaded from the HOSTS database (nhm.ac.uk/hosts; Robinson et al. 2015), a collection of published records of worldwide caterpillar host-plants. Hemipteran host-use data were downloaded from the Tri-Trophic Thematic Collection Network database (tcn.amnh.org), a compilation of insect collection records from academic museums in the United States. For both datasets, we restricted our analysis to records from North America (all localities labeled USA, Canada, Mexico or Nearctic). All plant taxonomic names were standardized with the Taxonomic Name Resolution Service (Boyle et al. 2013) and insect taxonomic names with the python package TaxonNamesResolver and the following reference databases: Aphid Species File (Favret 2015), Integrated Taxonomic Information System (itis.gov), and Catalogue of Life (catalogueoflife.org). We created binary presence/absence matrices of lepidopteran and hemipteran species by host plant order, with insects considered present on all hosts for which they had at least one host-use record. To focus computational resources on host taxa with enough statistical power to detect evolutionary host-use interactions, we restricted our analyses to focal host orders used by at least 100 insect species in one insect order (~10% of the total focal insect species per order).

We estimated time-scaled phylogenies for the North American lepidopteran and hemipteran species in our host-use dataset using a phyloinformatic approach (see Supplemental Materials for details). Phylogenetic data were not available for all species in the host-use dataset, but there was an overlap of host-use and phylogenetic data for 1604 lepidopteran species and 955 hemipteran species. Phylogenies and host-use matrices for these species are available on Dryad (datadryad.org).

**Statistical Analysis**. We used the insect-species-by-plant-order presence/absence data to investigate whether our focal host-use traits (presence/absence on each plant order) were positively or negatively correlated across the insect species. These correlations quantified whether insect species present on plant order A were more or less likely to be present on plant order B than expected by chance. For each insect order (Hemiptera and Lepidoptera) and each pairwise comparison between host-use traits, we set up a phylogenetic mixed model (Hadfield and Nakagawa 2010) with a logit link function (to accommodate binary data) using the package MCMCglmm (Hadfield 2010) in the R statistical environment (R Core Team 2015). We estimated both phylogenetic and residual correlations between the two host-use traits using the `“random=~us(trait): insect”` and `“rcov=~us(trait) : units”` syntax (Hadfield 2010). Prior parameter distributions were specified as `“prior<-list(R=list(V=diag(2),nu=2)`, `G=list(G1=list(V=diag(2),nu=2)))”`, and the mean of the posterior distribution was taken as the final estimate for each parameter. All MCMC chains ran for 10 million iterations with a burn-in of 1 million iterations, and we evaluated the convergence of ten chains for each model. Gelman-Rubin convergence analysis of each model's ten chains produced potential scale reduction factors under 1.10 in every case (96% were under 1.01), suggesting that all chains successfully converged (Gelman and Rubin 1992).

After separately estimating all pairwise evolutionary correlations between the focal host-use traits, we evaluated two emergent properties of the host-use network as a whole. First, we calculated the mean of all correlations involving each host-use trait to summarize whether presence on that host tended to be positively or negatively correlated with presence on all other hosts. Second, we asked whether the host-use traits could be grouped into clusters that had positive correlations within them and negative correlations between them. To identify the most strongly supported clusters, we used a distance matrix calculated from the pairwise correlations between host-use traits to produce a dendrogram of associations between the traits. Agglomerative hierarchical clustering was performed with the “complete” method of the hclust function in the R package fastcluster (Müllner 2013). After obtaining the dendrogram, we evaluated all possible cluster divisions produced by pruning the dendrogram at a single “level” (from broadest, with all host-use traits in a single cluster, to narrowest, with each host-use trait in its own cluster). The support for a given set of clusters was defined as the sum of all correlations between host-use traits in the same cluster minus the sum of all correlations between host-use traits in different clusters. Thus, positive correlations within clusters and negative correlations between clusters increased the support score, while negative correlations within clusters and positive correlations between clusters reduced the support score. The set of cluster divisions with the highest support score was chosen as the best characterization of network structure.

We tested the statistical significance of the resulting values by comparing them to those calculated for 100 null datasets. Each null dataset was generated by simulating independent Brownian motion of a continuous character for performance on each focal host order along the insect phylogenies, plus an equal amount of normally distributed residual variation in the performance values. We converted the resulting continuous host performance values to a binary host presence/absence character by assuming that only the insect species with the highest performance values for each host taxon were present on that host, with the threshold set by matching the number of species using that host in the empirical data (Felsenstein 2012). We then calculated all pairwise correlations between the host-use traits, mean correlations per host-use trait, and whole-network structure as we did for the empirical data. Empirical individual host-use trait mean correlations were considered statistically significant when their absolute values exceeded the maximum absolute values of any individual mean in 95% of null datasets. Empirical network structures was considered statistically significant when their support scores exceeded the support scores of 95% the null datasets.

## Results

**Host-use in Lepidoptera and Hemiptera**. We obtained North American host-use records and phylogenetic data for 1604 caterpillar species and 955 bug species (Fig. 2). Eleven host-plant orders met our prevalence cut-off of 100 species from one insect order, and each of them met the cut-off for both Hemiptera and Lepidoptera: Asterales, Caryophyllales, Ericales, Fabales, Fagales, Lamiales, Malpighiales, Pinales, Poales, Rosales, and Sapindales. Interactions with these focal host-plant orders accounted for 77% of all insect-species-by-plant-order interactions in the Lepidoptera dataset and 57% in the Hemiptera dataset.

**Fig. 2.**
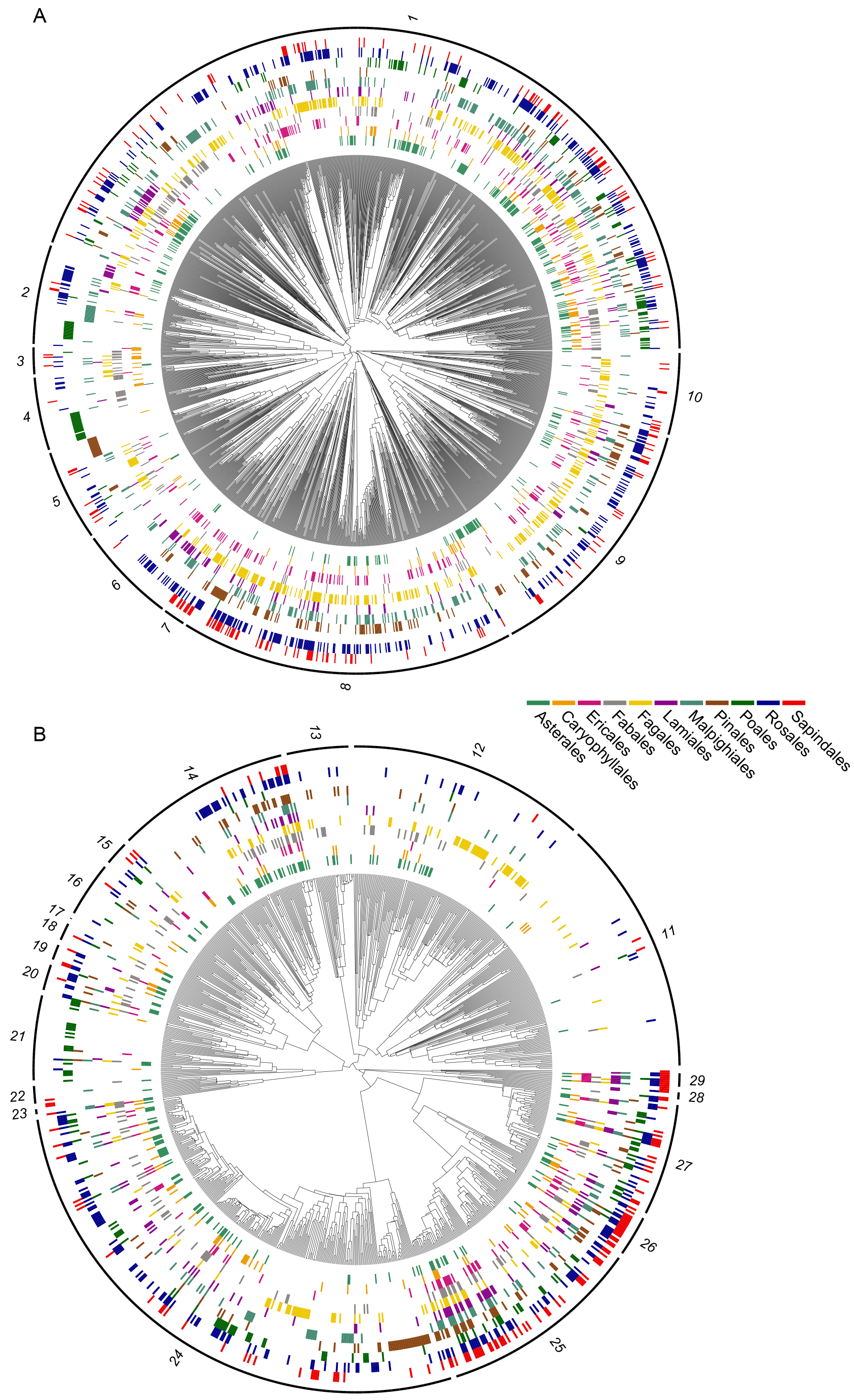
Maps of host-use traits on insect phylogenies. For each host-plant order, colored blocks indicate which insect species have been observed on that host. Insect species with no hosts shown were observed only on non-focal hosts or had no host-use information associated with their locality records (Hemiptera only). Insect families (and one superfamily) are indicated around the phylogenies as follows: (A) Lepidoptera – 1: Noctuoidea, 2: Nymphalidae, 3: Lycaenidae, 4: Hesperiidae, 5: Pyralidae, 6: Sphingidae, 7: Saturniidae, 8: Geometridae, 9: Tortricidae, 10: Gracillariidae. (B) Hemiptera – 11: Cicadellidae, 12: Membracidae, 13: Cicadidae, 14: Miridae, 15: Tingidae, 16: Pentatomidae, 17: Scutelleridae, 18: Coreidae, 19: Rhopalidae, 20: Lygaeidae, 21: Delphacidae, 22: Fulgoridae, 23: Flatidae, 24: Aphididae, 25: Diaspididae, 26: Coccidae, 27: Pseudococcidae, 28: Psylloidea, 29: Aleyrodidae.

**Host-use Correlations**. We recovered both positive and negative correlations between use of the focal host orders in the Lepidoptera, but mostly positive correlations in the Hemiptera (Fig. 3). The network of phylogenetic correlations between lepidopteran use of the focal host orders was significantly structured (*P* < 0.01), revealing two large clusters of host taxa (Fig. 4a). Cluster membership was phylogenetically diverse: the gymnosperm order Pinales (conifers) and monocot order Poales (grasses) were each affiliated with a different set of eudicot orders. Residual correlations between lepidopteran use of the focal host taxa also showed significant network structure (P < 0.01) but on this timescale use of all angiosperm hosts formed a single cluster of mostly positive associations (Fig. 4b). Use of Pinales was isolated from the angiosperm cluster, exhibiting a statistically significant negative mean pairwise correlation with use of the other hosts (−0.23, *P* < 0.01). In contrast, hempiteran host-use correlations indicated significant support for a single host-use cluster encompassing all focal hosts over both phylogenetic (P < 0.01, Fig. 4c) and residual timescales (P < 0.01, Fig. 4d).

**Fig. 3.**
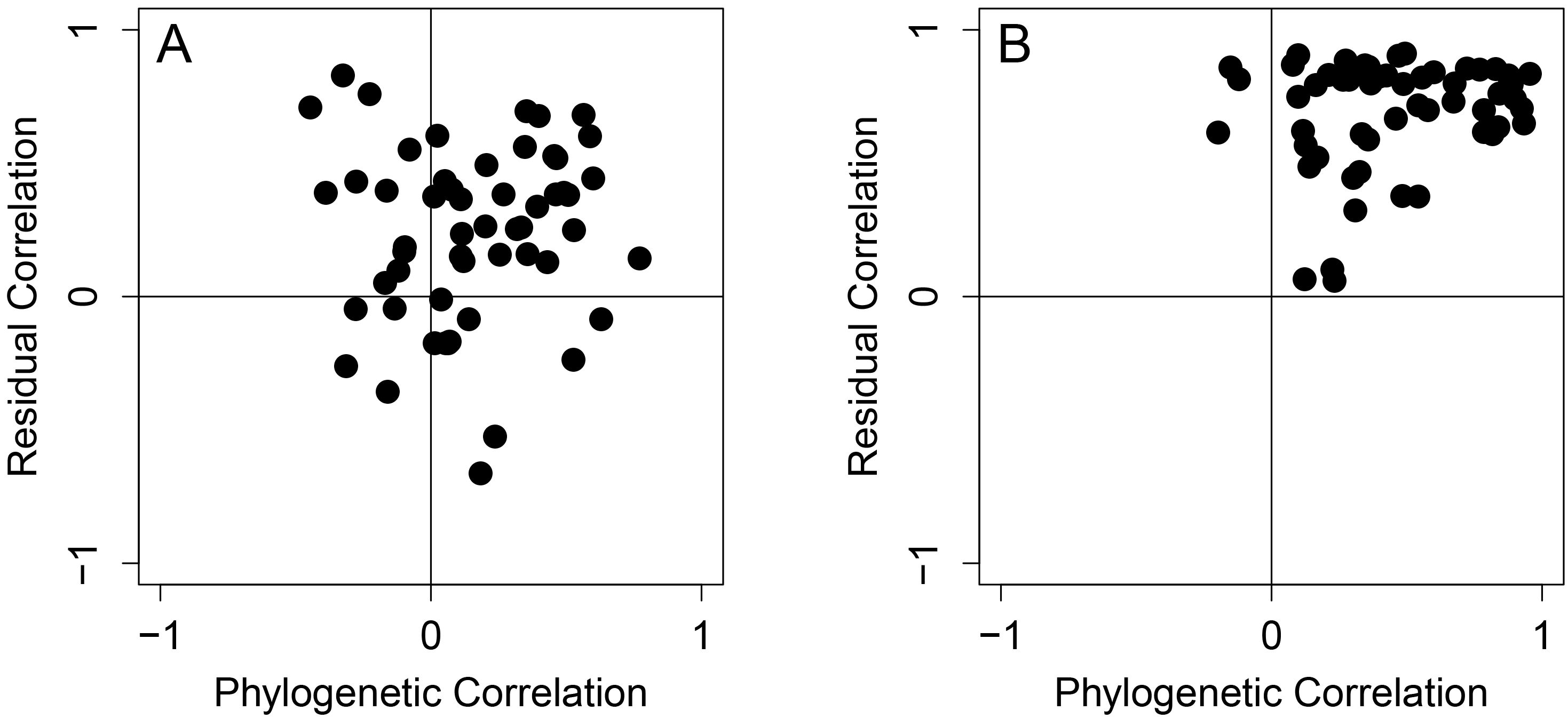
Empirical phylogenetic correlation by residual correlation plots of all 55 pairwise combinations of the focal host orders for Lepidoptera (A) and Hemiptera (B).

**Fig. 4.**
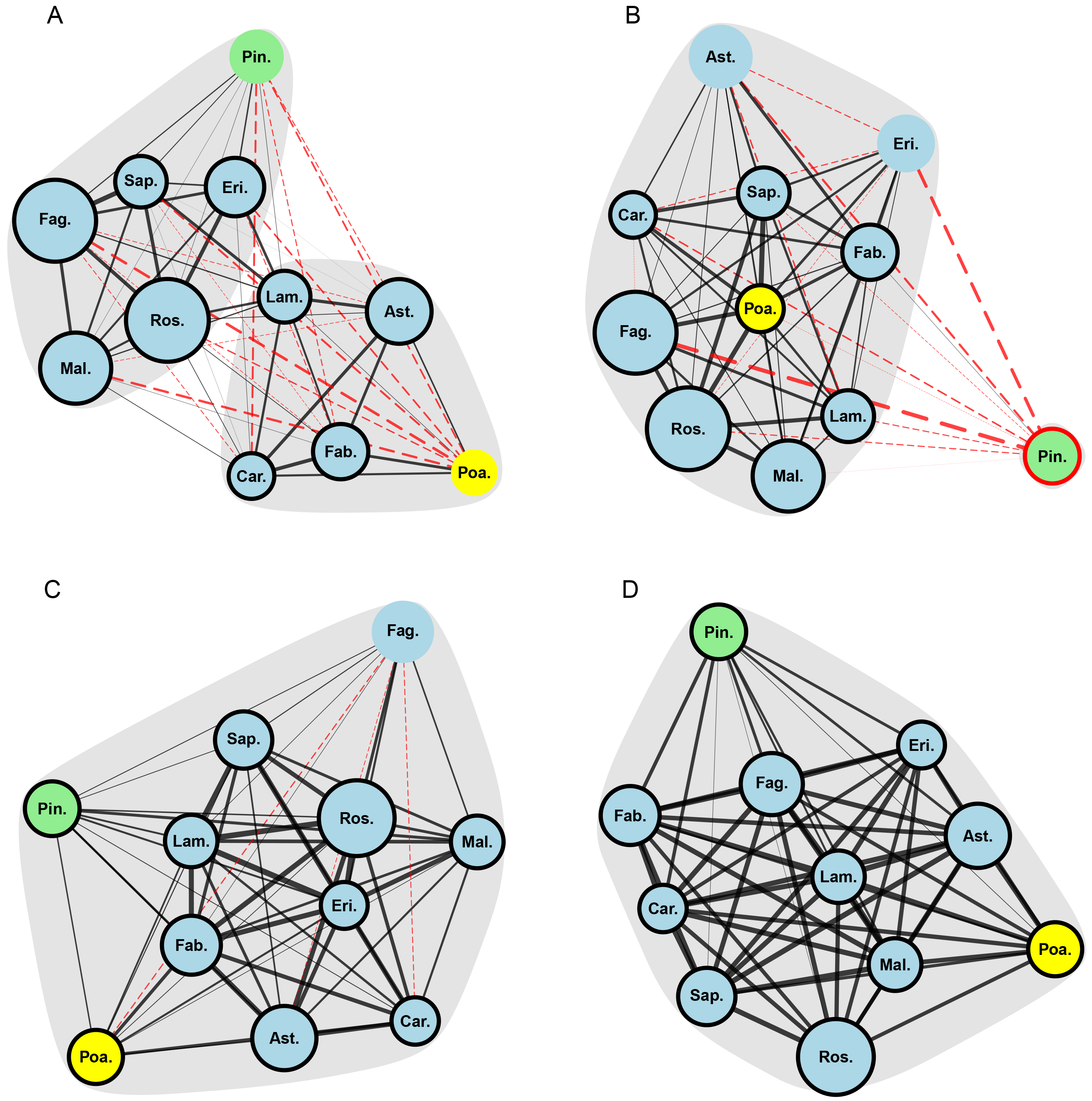
Network graphs of pairwise host-use trait correlations. (A) Lepidoptera - phylogenetic correlations. (B) Lepidoptera - residual correlations. (C) Hemiptera - phylogenetic correlations. (D) Hemiptera - residual correlations. Each vertex represents a host order, with vertex area proportional to the number of insects that were observed on that host. Positive interactions between presence on a pair of hosts are represented by solid, black lines and negative correlations by dashed, red lines, with line thickness proportional to the magnitude of the correlation. Network spatial structure was determined using the Kamada-Kawai (1989) algorithm, a force-directed layout method in which “repulsion” between vertices was proportional to the inverse of one plus the correlation values between the respective hosts. Vertices are labeled with the following abbreviations - Ast.: Asterales, Car.: Caryophyllales, Erl: Ericales, Fab.: Fabales, Fag.: Fagales, Lam.: Lamiales, Mal.: Malpighiales, Pin.: Pinales, Poa.: Poales, Ros.: Rosales, Sap.: Sapindales. Vertices are colored by taxonomic group - eudicots: blue, monocots: yellow, conifers: green. Statistically significant clusters (*P* < 0.05) are indicated by grey bubbles. Individual host orders with mean correlations of significantly higher magnitude than expected (*P* < 0.05) are indicated by bold vertex outlines (black for positive means, red for negative means).

## Discussion

Most models of the evolution of ecological specialization assume negative interactions (tradeoffs) between adaptations to different environments (Ravigné et al. 2009), but such interactions could also be neutral or positive (Gompert et al. 2015; Peterson et al. 2015). Here we used a statistical, phylogenetic approach to estimate the micro-and macroevolutionary correlations between use of eleven common host plant orders in both caterpillars and true bugs. Our results suggest that distinct micro-and macroevolutionary trade-offs constrain host-use in caterpillars, but use of all focal hosts are positively correlated on both timescales in true bugs. Overall, positive interactions between host-use adaptations appear to be more common than trade-offs in these plant-feeding insects.

We found some support for the idea that microevolutionary constraints (e.g. antagonistic pleiotropy) can produce host-use trade-offs in plant-feeding insects; lepidopteran presence on angiosperms was negatively correlated with presence on conifers over a short-term, phylogenetically independent timescale. This pattern suggests that caterpillar species tend to be found either angiosperm or conifer hosts (not both), yet they can shift between these alternative host-plant clades over relatively short evolutionary timescales. Such trade-offs between labile but mutually exclusive host-use traits are particularly significant because they can promote rapid speciation (Nosil et al. 2002) and adaptive radiations (Farrell 1998; Janz et al. 2006). In this case, microevolutionary constraints appear to reflect ancient phylogenetic divergence between host clades (Soltis et al. 2011). A similar pattern of microevolutionary trade-offs between use of phylogenetically distant hosts has been observed in networks of ecological interactions between fleas and their mammal hosts (Hadfield et al. 2014) and pollinators and their plant hosts (Rafferty and Ives 2013). Nevertheless, the prevalence of such constraints in plant-feeding insects, for instance between alternative host genera or species, remains unclear given that the single microevolutionary trade-off observed here occurred over the largest phylogenetic distance present among our focal host plant orders.

Most theoretical work on host-use evolution has focused on microevolutionary trade-offs, but we found that host-use constraints can also act over longer, macroevolutionary timescales. Over the phylogeny of the Lepidoptera, we observed a negative correlation between presence on hosts in two large, taxonomically diverse clusters. Interestingly, the clusters appeared to divided by morphology rather than phylogeny, with predominantly woody plant taxa (e.g. Pinales, Fagales) in one cluster and predominantly herbaceous taxa (e.g. Asterales, Poales) in the other. This pattern could reflect a longterm trade-off for lepidopteran lineages between use of alternative host growth forms or the habitats where those growth forms are found (Futuyma 1976). However, it is difficult to attribute macroevolutionary patterns to any particular mechanism. The phylogenetic correlations we detected here could be driven by any number of processes, including the accumulation of epistatic interactions (Weinreich et al. 2005; Remold 2012), dynamic evolution of host performance and host choice (Whitlock 1996; Ravigné et al. 2009), or geographic specificity of plant and insect lineages (Hadfield et al. 2014). Regardless, host-specificity in the Lepidoptera 1is clearly influenced by macroevolutionary processes that may be undetectable within a single insect population.

In contrast to the patterns observed in the Lepidoptera, hemipterans showed mostly positive associations between use of all focal host taxa over both micro-and macroevolutionary timescales. This surprising result suggests that generalist adaptations that increase fitness across multiple hosts have been more important for Hemiptera than specialist adaptations to particular hosts (Peterson et al. 2015). Moreover, hemipteran generalism appears completely unrestrained by host taxonomy even over very long timescales, leading to the evolution of both super-generalist species and clades where generalist strategies are common (Normark and Johnson 2011). However, we do not account for differences in fecundity between specialist and generalist insects on particular hosts; it may be that generalists usually have lower fitness - i.e. they are jacks of all trades but masters of none (Futuyma and Moreno 1988). Nevertheless, costs of generalism have been difficult to document (Forister et al. 2012; Gompert et al. 2015), so the positive residual correlations we observed may instead represent evolutionary breakthroughs made possible by novel mechanisms of phenotypic plasticity or other generalist adaptations (Barrett and Heil 2012).

There are many differences between Lepidoptera and Hemiptera, but their fundamentally distinct relationships with host plants may be important to understanding why evolutionary interactions between host-use traits appear to be different in the two groups (Pires and Guimarães 2012). Hemiptera are sucking insects, while Lepidoptera are generally leaf-chewers (Forister et al. 2015). These two feeding modes elicit different modes of plant defensive responses (Ali and Agrawal 2012), and sap-sucking may be particularly amenable to generalist adaptations that circumvent host defenses (Barrett and Heil 2012). In contrast, Lepidoptera often rely on specialized enzymes to detoxify defensive chemicals (Berenbaum and Feeny 2008), which may constrain the evolution of generalism, although generalist Lepidoptera do exist, possibly powered by phenotypic plasticity in enzyme expression (Yu et al. 1979; Li et al. 2002).

Overall, the relatively few, broad-scale trade-offs found here fail to explain the prevalence of specialization in plant-feeding insects, which are often restricted to hosts in a single plant family or genus (Forister et al. 2015). Our analysis grouped hosts by order, obscuring potential variation within orders in defensive strategies; host plant families or genera with strong or physiologically unique defenses may be more likely to produce trade-offs for plant-feeding insects than host plant taxa with weaker or more common defenses. However, our previous study of host-use in the large hemipteran family Diaspididae found positive correlations between use of all hosts but one within a network of 64 taxonomically diverse host genera (Peterson et al. 2015), indicating that greater taxonomic resolution does not necessarily reveal trade-offs between host-use traits. We also took a broad approach in looking for correlations between host-use traits across whole insect orders, thereby overlooking any idiosyncratic trade-offs that may arise from the unique natural history of individual insect species. Species-specific trade-offs have been documented (e.g. Nosil et al. 2002), yet our results suggest that few microevolutionary trade-offs constrain host-use across large numbers of insect species. Thus, although trade-offs may emerge at any time due to novel epistatic interactions (Remold 2012; Satterwhite and Cooper 2015), the fact that generalist species frequently escape such trade-offs suggests that long-term evolutionary interactions between host-use traits are dominated by positively pleiotropic or neutral adaptations.

Trade-offs play an intuitive and possibly inescapable role in constraining performance across multiple tasks (Shoval et al. 2012), but performance limits may not define the ecological niches of plant-feeding insects. Alternative factors unrelated to differential growth or survival on alternative hosts, such as mate-finding (Hawthorne and Via 2002) or neural constraints in host identification (Bernays 2001), may shape the evolution of each species' ecological niche. Host-range may also be limited by genetic drift even if adaptive interactions between host-use traits are positive or neutral (Gompert et al. 2015). Specialization-by-drift might be particularly significant in a geographical context, as interactions between host-range and geographical range can strongly affect the host-use selection pressures experienced by an insect lineage (Janz and Nylin 2008). In the absence of much evidence for negative interations between host-use adaptations in plant-feeding insects, a neutral model may be the best default both for the structure ecological networks (Canard et al. 2014) and how those networks evolve over time.

**Supplementary Information** is available in the online version of the paper.

## Acknowledgements

We thank D. Moen, A. Porter, L. Doubleday, S. Noda and three anonymous reviewers for comments that improved the manuscript. This work was supported by the National Science Foundation (EF -1115191 and DEB-1258001)

## Author Contributions

All authors were involved in the study design and wrote the manuscript. D.A.P. and N.B.H. analyzed the data.

## Author Information

The authors declare no competing financial interests. Correspondence and requests for materials should be addressed to D.A.P.

## Literature Cited

Agrawal, A. A., J. K. Conner, and S. Rasmann. 2010. Tradeoffs and negative correlations in evolutionary ecology. Pages 243–268 in Evolution since Darwin: The first 150 years.

Ali, J. G., and A. A. Agrawal. 2012. Specialist versus generalist insect herbivores and plant defense. Trends in Plant Science 17:293–302.

Barrett, L. G., and M. Heil. 2012. Unifying concepts and mechanisms in the specificity of plant-enemy interactions. Trends in Plant Science 17:282–292.

Berenbaum, M. R., and P. P. Feeny. 2008. Chemical mediation of host-plant specialization: the papilionid paradigm. Pages 3–19 in Specialization, Speciation, and Radiation: The Evolutionary Biology of Herbivorous Insects.

Bernays, E. A. 2001. Neural limitations in phytophagous insects: implications for diet breadth and evolution of host affiliation. Annual Review of Entomology 46:703–727.

Boyle, B., N. Hopkins, Z. Lu, J. A. Raygoza Garay, D. Mozzherin, T. Rees, N. Matasci, et al. 2013. The taxonomic name resolution service: an online tool for automated standardization of plant names. BMC Bioinformatics 14:16.

Campbell, S. A., and A. Kessler. 2013. Plant mating system transitions drive the macroevolution of defense strategies. proceedings of the National Academy of Sciences 110:3973–3978.

Canard, E. F., N. Mouquet, D. Mouillot, M. Stanko, D. Miklisova, and D. Gravel. 2014. Empirical evaluation of neutral interactions in host-parasite networks. The American Naturalist 183:468–79.

Elena, S. F., and R. E. Lenski. 2003. Evolution experiments with microorganisms: the dynamics and genetic bases of adaptation. Nature reviews. Genetics 4:457–69.

Farrell, B. D. 1998. “Inordinate fondness” explained: why are there so many beetles? Science 281:555–559.

Favret, C. 2015. Aphid species file. [aphid.speciesfile.org]

Felsenstein, J. 2012. A comparative method for both discrete and continuous characters using the threshold model. The American Naturalist 179:145–156.

Forister, M. L., L. A. Dyer, M. Singer, J. O. Stireman, and J. T. Lill. 2012. Revisiting the evolution of ecological interactions. Ecology 93:981–991.

Forister, M. L., V. Novotny, A. K. Panorska, L. Baje, Y. Basset, P. T. Butterill, L. Cizek, et al. 2015. The global distribution of diet breadth in insect herbivores. Proceedings of the National Academy of Sciences of the United States of America 112:442–7.

Fry, J. 1996. The evolution of host specialization: are trade-offs overrated? The American Naturalist 148:S84–107.

Futuyma, D. J. 1976. Food plant specialization and environmental predictability in Lepidoptera. The American Naturalist 110:285–292.

Futuyma, D. J.. 2008. Sympatric speciation: norm or exception? Pages 136–147 in Specialization, Speciation, and Radiation: The Evolutionary Biology of Herbivorous Insects.

Futuyma, D. J., and A. A. Agrawal. 2009. Macroevolution and the biological diversity of plants and herbivores. Proceedings of the National Academy of Sciences of the United States of America 106:18054–18061.

Futuyma, D. J., M. Keese, and D. J. Funk. 1995. Genetic constraints on macroevolution: the evolution of host affiliation in the leaf beetle genus Ophraella. Evolution 49:797–809.

Futuyma, D. J., and G. Moreno. 1988. The evolution of ecological specialization. Annual Review of Ecology and Systematics 19:207–233.

Gelman, A., and D. B. Rubin. 1992. Inference from iterative simulation using multiple sequences. Statistical Science 7:457–472.

Gompert, Z., J. P. Jahner, C. F. Scholl, J. S. Wilson, L. K. Lucas, V. Soria-Carrasco, J. A. Fordyce, et al. 2015. The evolution of novel host use is unlikely to be constrained by trade-offs or a lack of genetic variation. Molecular Ecology 24:2777–2793.

Hadfield, J. D. 2010. MCMC methods for multi-response generalized linear mixed models: The MCMCglmm R package. Journal of Statistical Software 33:1–22.

Hadfield, J. D., B. R. Krasnov, R. Poulin, and S. Nakagawa. 2014. A tale of two phylogenies: comparative analyses of ecological interactions. The American Naturalist 183:174–87.

Hadfield, J. D., and S. Nakagawa. 2010. General quantitative genetic methods for comparative biology: Phylogenies, taxonomies and multi-trait models for continuous and categorical characters. Journal of Evolutionary Biology 23:494–508.

Hawthorne, D. J., and S. Via. 2002. The genetic architecture of ecological specialization: correlated gene effects on host use and habitat choice in pea aphids. The American Naturalist 159:S76–S88.

Janz, N., and S. Nylin. 2008. The oscillation hypothesis of host-plant range and speciation. Pages 203–215 in Specialization, Speciation, and Radiation: The Evolutionary Biology of Herbivorous Insects.

Janz, N., S. Nylin, and N. Wahlberg. 2006. Diversity begets diversity: host expansions and the diversification of plant-feeding insects. BMC Evolutionary Biology 6:4.

Johnson, M. T. J., A. R. Ives, J. Ahern, and J. P. Salminen. 2014. Macroevolution of plant defenses against herbivores in the evening primroses. New Phytologist 203:267–279.

Joshi, A., and J. N. Thompson. 1995. Trade-offs and the evolution of host specialization. Evolutionary Ecology 9:82–92.

Kamada, T., and S. Kawai. 1989. An algorithm for drawing general undirected graphs. Information Processing Letters 31:7–15.

Li, W., M. A. Schuler, and M. R. Berenbaum. 2003. Diversification of furanocoumarin-metabolizing cytochrome P450 monooxygenases in two papilionids: specificity and substrate encounter rate. Proceedings of the National Academy of Sciences of the United States of America 100:14593–14598.

Li, X., M. A. Schuler, and M. R. Berenbaum. 2002. Jasmonate and salicylate induce expression of herbivore cytochrome P450 genes. Nature 419:712–715.

Maddison, W. P., and R. G. FitzJohn. 2015. The unsolved challenge to phylogenetic correlation tests for categorical characters. Systematic Biology 64:127–136.

Müllner, D. 2013. fastcluster: fast hierarchical, agglomerative clustering routines for R and Python. Journal of Statistical Software 53:1–18.

Normark, B. B., and N. A. Johnson. 2011. Niche explosion. Genetica 139:551–564.

Nosil, P., B. J. Crespi, and C. P. Sandoval. 2002. Host-plant adaptation drives the parallel evolution of reproductive isolation. Nature 417:440–3.

Nurmi, T., and K. Parvinen. 2011. Joint evolution of specialization and dispersal in structured metapopulations. Journal of Theoretical Biology 275:78–92.

Peterson, D. A., N. B. Hardy, G. E. Morse, I. C. Stocks, A. Okusu, and B. B. Normark. 2015. Phylogenetic analysis reveals positive correlations between adaptations to diverse hosts in a group of pathogen-like herbivores. Evolution 69:2785–2792.

Pires, M. M., and P. R. Guimarães. 2012. Interaction intimacy organizes networks of antagonistic interactions in different ways. Journal of The Royal Society Interface rsif20120649.

R Core Team. 2015. R: A language and environment for statistical computing. R Foundation for Statistical Computing, Vienna, Austria.

Rafferty, N. E., and A. R. Ives. 2013. Phylogenetic trait-based analyses of ecological networks. Ecology 94:2321–2333.

Ravigné, V., U. Dieckmann, and I. Olivieri. 2009. Live where you thrive: joint evolution of habitat choice and local adaptation facilitates specialization and promotes diversity. The American Naturalist 174:E141–E169.

Remold, S. 2012. Understanding specialism when the jack of all trades can be the master of all. Proceedings of the Royal Society B: Biological Sciences 279:4861–4869.

Robinson, G. S., P. R. Ackery, I. J. Kitching, G. W. Beccaloni, and L. M. Hernández. 2015. HOSTS - a database of the world's Lepidopteran hostplants. Natural History Museum, London.

Rodriguez-Verdugo, A., D. Carrillo-Cisneros, A. Gonzalez-Gonzalez, B. S. Gaut, and A. F. Bennett. 2014. Different tradeoffs result from alternate genetic adaptations to a common environment. Proceedings of the National Academy of Sciences 111:12121–12126.

Satterwhite, R. S., and T. F. Cooper. 2015. Constraints on adaptation of Escherichia coli to mixed-resource environments increase over time. Evolution 69:2067–2078.

Scheirs, J., K. Jordaens, and L. De Bruyn. 2005. Have genetic trade-offs in host use been overlooked in arthropods? Evolutionary Ecology 19:551–561.

Scriber, J. M. 2010. Integrating ancient patterns and current dynamics of insect-plant interactions: Taxonomic and geographic variation in herbivore specialization. Insect Science 17:471–507.

Shoval, O., H. Sheftel, G. Shinar, Y. Hart, O. Ramote, A. Mayo, E. Dekel, et al. 2012. Evolutionary trade-offs, Pareto optimality, and the geometry of phenotype space. Science 336:1157–1160.

Soltis, D. E., S. A. Smith, N. Cellinese, K. J. Wurdack, D. C. Tank, S. F. Brockington, N. F. Refulio-Rodriguez, et al. 2011. Angiosperm phylogeny: 17 genes, 640 taxa. American Journal of Botany 98:704–30.

Weinreich, D. M., R. A. Watson, and L. Chao. 2005. Perspective: sign epistasis and genetic constraint on evolutionary trajectories. Evolution 59:1165–1174.

Whitlock, M. C. 1996. The red queen beats the jack-of-all-trades: the limitations on the evolution of phenotypic plasticity and niche breadth. American Naturalist 148:S65–S77.

Yu, S. J., R. E. Berry, and L. C. Terriere. 1979. Host plant stimulation of detoxifying enzymes in a phytophagous insect. Pesticide Biochemistry and Physiology 12:280–284.

